# Prophage acquisition by *Staphylococcus aureus* contributes to the expansion of Staphylococcal immune evasion

**DOI:** 10.1101/2023.04.27.538627

**Authors:** Roshan Nepal, Ghais Houtak, George Bouras, Mahnaz Ramezanpour, Sholeh Feizi, Gohar Shaghayegh, Keith Shearwin, Alkis James Psaltis, Peter-John Wormald, Sarah Vreugde

## Abstract

*Staphylococcus aureus* colonizes 30% of the human population, but only a few clones cause severe infections. *S. aureus’* virulence varies and partly depends on the presence of prophages, viral DNA embedded in the *S. aureus* core genome, such as hlb-converting prophage (ϕSa3int). Human-adapted *S. aureus* often harbours a ϕSa3int group of prophages preferentially integrated into their β-hemolysin (*hlb*) gene that encodes human immune evasion cluster (IEC) genes. Exotoxins and immune modulatory molecules encoded by this prophage can inhibit human innate immunity increasing *S. aureus* pathogenicity. This study aims to investigate the genomic and phenotypic plasticity of *S. aureus* and changes in its extracellular proteome after the acquisition of ϕSa3int prophage.

To achieve this, we used *S. aureus* strains isolated from the sinus cavities of a patient with severe chronic rhinosinusitis (CRS) at two different time points (*S. aureus* SA222 and *S. aureus* SA333) and hybrid sequenced the strains using short-read Illumina and long-read Oxford nanopore technology. *In silico* analysis showed the presence of a ϕSa3int prophage in the later isolate but not in the earlier isolate while most of the core genes remained identical. Using mitomycin C, we induced the ϕSa3int prophage, and transduced it into the Sa3int-prophage-free SA222 isolate to obtain a laboratory generated ‘double lysogen’. We confirmed the successful lysogenisation with culture methods (spot assay, blood agar) and also by sequencing. We compared growth kinetics, biofilm biomass and metabolic activity between parent and the lysogen by establishing growth curves, crystal violet and resazurin assays. Exoproteins were identified and quantified using mass spectrophotometry.

Integration of ϕSa3int prophage in SA222 down-regulated the beta-hemolysin expression of the lysogen*. In silico* analysis of the *S. aureus* genome confirmed the insertion of a ∼43.8 kb ϕSa3int prophage into *hlb* gene. Insertion of prophage DNA did not alter the growth kinetics, biofilm formation, adhesion to primary human nasal epithelial cells and the metabolic activity in a biofilm. However, the acquisition of ϕSa3int prophage significantly changed the expression of various secreted proteins, both bacterial and prophage-encoded. Altogether, thirty-eight exoproteins were significantly differentially regulated in the laboratory created lysogen, compared to its recipient strain SA222. Among these proteins, there was significant upregulation of 21 exoproteins (55.3 %) including staphylokinase (sak), SCIN (scn), and intercellular adhesion protein B (icaB) and downregulation of 17 exoproteins (44.7 %), including β-hemolysin (hlb/sph) and outer membrane porin (phoE). Most of the upregulated proteins were involved in immunomodulation that help *S. aureus* escape human innate immunity and help cause chronic infection. These findings may contribute to the development of novel approaches to render S. *aureus* susceptible to the immune response by blocking prophage-associated defence mechanisms.

**Highlights:**

- A ϕSa3int prophage preferentially integrates into the β-haemolysin gene (*hlb*) gene thereby disrupting the beta-hemolysin function.
- A ∼43.8 kb ϕSa3int prophage acquisition by *S. aureus* has no impact on its growth kinetics, biofilm formation and adhesion to primary human nasal epithelial cells (HNECs).
- The presence of a ϕSa3int group prophage likely enhances *Staphylococcus aureus’* human immune evasion capability as the prophage encodes a complete set of immune evasion cluster (IEC) genes.
- Targeted identification of virulence factors in addition to species and strain identification may lead to better-personalized therapy as not all *S. aureus* carry the same virulence genes.

## 1. INTRODUCTION

Bacteria often harbour dormant phage DNA (prophages) embedded within their chromosome. These prophage sequences can confer auxiliary functions, frequently increasing bacterial fitness. However, prophages can also carry various virulence factors (VF), toxins and antimicrobial resistance genes (ARGs), such that lysogens (bacteria carrying a prophage) are often considered more virulent than the corresponding prophage-free strain (Kondo et al. 2021, Davies et al. 2016). Prophages can be induced spontaneously or in response to various extrinsic factors like UV, sub-lethal antibiotics and chemicals (Henrot and Petit 2022). Although spontaneous prophage induction (SPI) usually occurs at a low level and kills a small fraction of bacterial cells within a population, the released phage particles lysogenize other susceptible strains in the niche, thereby transducing the prophage-encoded virulence factors (PVFs) horizontally. Further, the induction of prophages into phage particles can also activate an anti-viral immune response in mammalian cells that protects bacteria from phagocytosis (Gogokhia et al. 2019, Popescu et al. 2021). It is well established that prophage induction is enhanced under sub-lethal concentrations of various antibiotics and chemicals (Allen et al. 2011, Boling et al. 2020, Goerke, Koller and Wolz 2006). Moreover, some studies have also found that prophage domestication and induction enhance biofilm formation, further increasing bacterial survival and fitness (Nanda, Thormann and Frunzke 2015, Carrolo et al. 2010). As bacteria often acquire virulent traits under inadequate or inappropriate antimicrobial treatment regimens despite the DNA replication cost involved, it is essential to understand prophage diversity, prophage dispersal and their role in virulence dissemination. Most bacterial infections are caused by a small number of successful clones that are virulent and pathogenic compared to the commensal (Beceiro, Tomás and Bou 2013). Therefore, understanding the dynamics of prophage-mediated dissemination of prophage-encoded virulence may provide information about the origin and spread of virulent clones.

*Staphylococcus aureus* is a genetically and metabolically diverse, highly successful opportunistic bacterial pathogen colonizing the mucosal surfaces of approximately 30 % of humans (Tong et al. 2015)*. S. aureus* is often isolated from the sinuses of chronic rhinosinusitis (CRS) patients, more often in CRS with nasal polyposis (CRSwNP) (Vickery, Ramakrishnan and Suh 2019). Factors related to virulence in *S. aureus* are often associated with mobile genetic elements (MGEs) like plasmids, insertion sequences and prophages, suggesting that horizontal acquisition of MGEs plays a significant role in the development of the virulence (Howden et al. 2023). A well-known example is β-hemolysin-converting prophages (hereafter ϕSa3int) carrying various immune evasion cluster (IEC) genes (*sak, chp, scn* and *sea/sep)* that protect bacteria from neutrophil-dependent phagocytosis (van Wamel et al. 2006, Nepal et al. 2022). Earlier, we reported that *S. aureus* isolated from CRSwNP patients often harboured ϕSa3int prophages that disrupt the production of β-hemolysin (Nepal et al. 2021). β-hemolysin is a sphingomyelinase hemolysin that significantly contributes to *S. aureus* pathogenesis (Tran et al. 2019), reduces the ciliary activity of nasal epithelial cells and induces sinusitis (Kim et al. 2000). However, the pathogenesis of β- hemolysin toxin in humans is argued due to negative conversion by the ϕSa3int prophage which is present in the majority of *S. aureus* colonising humans (Salgado-Pabon et al. 2014). Widespread distribution ϕSa3int prophages among *S. aureus* isolated from the nasal cavity of humans suggests that the induction and re-integration of the released phages drive the dissemination of virulence genes, particularly IEC, contributing to the genetic diversification and functional adaptations of *S. aureus* (Chaguza et al. 2022). However, the presence of prophage DNA does not imply the functionality of the virulence genes it carries.

This study aimed to investigate ϕSa3int prophage-mediated phenotypic alteration and virulence expression in *S. aureus* isolated from CRS patients. By transducing an intact ϕSa3int prophage, induced from *S. aureus* (SA333) isolated from a severe CRS patient, into another genetically close ϕSa3int-free clinical strain isolated from the same patient at a different time point (SA222), we demonstrated that ϕSa3int prophages integrate into *hlb* gene and significantly upregulates expression of various exotoxins responsible for human immune evasion. However, the domestication of the prophage did not alter the lysogen’s growth kinetics and biofilm properties. Hence, ϕSa3int prophages are crucial factors in the dissemination of IEC genes and virulence in *S. aureus* that may contribute to the chronic colonization in the nasal cavity of CRS patients.

## 2. MATERIALS AND METHODS

### 2.1 Ethics, bacterial strains, cells and growth conditions

Ethics approval for the use of clinical isolates (CIs) and primary human nasal epithelial cells (HNECs) was obtained from the Human Research Ethics Committee of the Central Adelaide Local Health Network (HREC/18/CALHN/69). All the *S. aureus* clinical isolates used in this study were retrieved from glycerol stocks and cultured at 37°C overnight on nutrient agar (NA, Oxoid Ltd, Hampshire, UK). *S. aureus* RN4220 and *S. aureus* ATCC25923 were from the German Collection of Microorganisms and Cell Cultures (DSMZ, GmbH) and American Type Culture Collection (ATCC, Manassas, USA) respectively. An isolated colony was propagated in 15.0 ml tryptic soy broth (TSB, 1X, Oxoid Ltd, Hampshire, UK) overnight in a shaking incubator (180 rpm) at 37°C unless stated otherwise. The HNECs used for the adhesion assay were collected from a non-CRS (control) patient at the time of surgery.

### 2.2 Genomic DNA extraction, sequencing and genome assembly

The genomic DNA (gDNA) of all *S. aureus* CIs were extracted using DNeasy Blood & Tissue Kit (Cat. #69504, Qiagen Pty. Ltd, Australia) according to the manufacturer’s guidelines with slight modifications. Briefly, 700 µl of overnight broth culture in TSB was centrifuged (4000 x g) in a 1.5 ml Eppendorf tube for 10 minutes. The pellet was suspended in 180 µl of enzymatic lysis buffer (20 mM Tris-Cl, pH8; 2mM sodium EDTA; 1.2% Triton X-100, 200 µg/ml final concentration lysostaphin, filter sterilized) and incubated at 37°C for 30 minutes. Then 25 μl of proteinase K and 200 μl of Buffer AL (undiluted, provided with extraction kits) were added and mixed by vortexing followed by incubation at 56°C. After 30 minutes, 200 μl of 99% ethanol (chilled) was added and mixed by vortexing. The mixture was then transferred to DNeasy Mini Spin column (Qiagen Pty. Ltd, Australia, Cat. #69504), and DNA was extracted following the manufacturer’s guidelines.

The extracted gDNA was sequenced using the short-read Illumina platform (Illumina Inc, San Diego, USA) and in-house long-read Oxford Nanopore Technology (ONT) using the MinION Mk1C device (Oxford Nanopore Technologies, Oxford, UK) following the manufacturer’s instructions and in-house established protocol (Shaghayegh et al. 2023). Briefly, the short-read sequencing was done on Illumina NextSeq 550 platform using NextSeq 500/550 Mid-Output kit v2.5 (Illumina Inc, San Diego, USA) at a commercial sequencing facility (SA Pathology, Adelaide, SA, Australia). Briefly, gDNA was isolated using the NucleoSpin Microbial DNA kit (Machery-Nagel GmbH, Duren, Germany). Sequencing libraries were prepared using a modified protocol for the Nextera XT DNA library preparation kit (Illumina Inc, San Diego, USA). The gDNA was fragmented and amplified using a low-cycle PCR reaction. After the manual purification and normalisation of the amplicon library, 150 bp reads were obtained. Long-read whole genome sequencing was performed using MinION flowcells (R9.4.1) with the Rapid Barcoding Kit (Oxford Nanopore Technology, UK, #Cat: SQK-RBK 110.96) according to the manufacturer’s instructions. In brief, 50 ng of gDNA from each CIs was used for sequencing. Base-calling was conducted with Guppy v6.2.11 (mode = super accuracy) using the ’dna_r9.4.1_450bps_sup.cfg’ configuration (Oxford Nanopore Technology, UK).

Complete chromosomal *S. aureus* assemblies were created using a customised Snakemake pipeline (Mölder et al. 2021) that is available at https://github.com/gbouras13/hybracter via the Snaketool (Roach et al. 2022) powered command line tool hybracter following the protocols outlined in Wick, Judd and Holt (2023). The complete pipeline can be found as supplementary text ST1. Chromosome assemblies were annotated with Bakta v1.6.1 (Schwengers et al. 2021a). All CIs were typed to determine sequence type (ST) and clonal complex (CC) according to the PubMLST database using MLST (Seemann mlst, Jolley, Bray and Maiden 2018). We used Snippy v4.6.0 to detect single nucleotide polymorphisms (SNPs) between CIs (Seemann 2015) and Abricate v1.0.1 to screen for anti-microbial resistance and virulence factor genes (Seemann 2020).

### 2.3 *In silico* identification of prophage and prophage annotation

Prophage regions in both *S. aureus* CIs (SA222 and SA333, isolated from the same patient at different time points) were first identified using PHASTEST (https://phastest.ca) and PhiSpy (https://github.com/linsalrob/PhiSpy) with default settings (Arndt et al. 2016, Akhter, Aziz and Edwards 2012, Wishart 2023). The exact genome of the ϕSa3int prophage was then curated with our in-house program hlbroken (https://github.com/gbouras13/hlbroken), which extracts the sequences between *hlb* gene only. The identified φSa2int and φSa3int prophage sequences were also annotated and visualized with Pharokka v1.2.0 (Bouras et al. 2023).

### 2.4 induction and multi-host range assay of induced prophages

Prophages from both CIs were induced using mitomycin C (MMC), purified and spotted on previously studied *S. aureus* CIs (N=66) using the soft-agar overlay technique as described earlier (Nepal et al. 2023) to study its multiple-host range. Briefly, MMC (final concentration = 1.0 μg/ml) (Sigma-Aldrich, Missouri, USA, #Lot: SLBX4310) was added to exponentially growing cells (OD_600_ = 0.3) in TSB and further incubated for 6 hrs at 37°C in a rotating incubator (180 rpm). Optical density at 600 nm (OD_600_) was measured every hour. After 6 hrs, the culture was centrifuged at 4000 rpm for 15 minutes at 4°C and the supernatant was filtered through 0.2 μm pore size syringe filter (13 mm, Acordisc^®^, Pall International, Fribourg, Switzerland, #Cat: 4612) to obtain pure phage lysate. Briefly, host bacteria were overlayed in double-layered TSA and 10.0 μl of purified lysate was spotted on the top agar in triplicates.

### 2.5 Construction and verification of lysogens

Ten microliters of purified phage lysate induced from SA333 were spotted on a soft agar overlaid with recipient bacteria SA222. The plates were dried and incubated overnight at 37°C. The next day, a loopful of bacteria from the lysis spots were streaked on sheep blood agar (SBA, ThermoFisher, Australia, #Cat: R01202) and incubated overnight at 37°C. Colonies without beta-hemolysis (possibly lysogens) were picked and sub-cultured in TSA. The stability of these constructs possibly harboring both ϕSa2int and ϕSa3int prophages was confirmed through multiple sub-cultures in SBA. Two constructs devoid of beta-hemolysin (hereafter SA-L1 and SA-L2) were picked for further analysis. The integrity and re-inducibility of integrated prophage were then confirmed by re-induction from the constructs using MMC as described earlier and spot assayed on SA222, SA333 and RN4220. The integration of ϕSa3int prophage into the *hlb* gene was verified by inspecting the *hlb* gene in assembled genomes using the Integrative Genomics Viewer (IGV) (Robinson et al. 2011).

### 2.6 Growth curve, biofilm biomass and biofilm metabolic activity

Bacterial growth kinetics were determined by measuring the optical density of broth culture at 600 nm (OD_600_). Briefly, 100 μl of 1.0 McFarland standard unit (MFU in saline, prepared from overnight cultured colonies on NA plates) was added to 15.0 ml of TSB in a 50 ml Falcon tube. The tubes were incubated at 37°C in a shaking incubator (180 rpm). Every hour, 100 µl of culture was removed and mixed with 900 µl of sterile TSB in a cuvette. The OD_600_ was then measured using a SmartSpec^TM^ 3000 UV/Vis spectrophotometer (Bio-Rad Laboratories Inc, California, USA).

The biofilm variation between the clinical isolates and lysogens was qualitatively assessed by culturing the bacteria on modified Congo red agar (CRA) (37 gm/l brain heart infusion broth supplemented with 50 g/l sucrose, 0.8 g/l Congo red stain and 1.0 % agar) according to Freeman et al. (Freeman, Falkiner and Keane 1989). Colony morphology of SA222, SA333 and SA-L1/SA-L2 on CRA was assessed after 48 hours of incubation at 37°C. Further, biofilm biomass and biofilm metabolic activity (cell viability) was performed using crystal violet (CV) assay and alamarBlue^®^ cell viability assay as per manufacturer’s guidelines (Life Technologies, Oregon, USA) in biofilms established for 48 hours in 96-well flat-bottomed (Costar, Corning Incorporated, USA. #Ref: 3599) and 96-well flat and clear bottom black assay plate (Costar, Corning Incorporated, USA. #Ref: 3603) respectively, as described earlier (Nepal et al. 2023). Briefly, the overnight NA culture of *S. aureus* was adjusted to 1.0 McFarland standard and diluted 1:15 in tryptic soy broth. One-hundred-fifty microliters of each diluted culture were pipetted into the inner wells of the respective 96-well plates. The peripheral wells were filled with sterile water, sealed with aluminium foil and incubated at 37°C in an orbital shaker (80 rpm). After 48 hours of incubation, planktonic cells were carefully aspirated, and the plates were washed twice with phosphate buffered saline (PBS, 1X) followed by the crystal violet or alamarBlue^®^ assay. For CV, plates were air-dried and 180 μl of 0.01% CV solution was added to each inner well and left at room temperature for staining. After 10 minutes, the excess CV was aspirated, washed twice with PBS (1X) and air-dried. Finally, the biomass-bound crystal violet was solubilized in 200 μl of 30% acetic acid. The biomass was measured in terms of absorbance (OD_600_) at 600 nm using CLARIOstar Plus (BMG Labtech, Ortenberg, Germany). Similarly, for biofilm metabolic activity, 200 μl working solution of alamarBlue^®^ (1X) was added to each inner well, covered with aluminium foil and incubated at 37°C in an orbital shaker (80 rpm). Resorufin fluorescence was monitored at the 1- hr interval for 6 hours using the CLARIOstar Plus (BMG Labtech, Ortenberg, Germany) microplate reader (excitation = 530 nm, emission = 590 nm). The metabolic activity (expressed as fluorescence units) at a 2-hour timepoint was considered for further analysis. All experiments were performed in triplicate with six technical replicates.

### 2.7 Adhesion assay of *S. aureus* to primary human nasal epithelial cells

The adhesion of *S. aureus* clinical strains and lysogens to primary human nasal epithelial cells (HNEC) was studied following the protocol by Yang and Ji with slight modifications (Yang and Ji 2014). Briefly, primary HNECs were cultured in RPMI 1640 working media (supplemented with 10% FBS and 1% antibiotic-antimycotic, hereafter RPMI 1640-WM) to 70% confluency in a tissue culture flask (T-75, Sarstedt, Nümbrecht, Germany) at 37°C, 5% CO_2_ incubator and transferred (1.0 ml/well) into a 24-well tissue culture plate (Sarstedt, Nümbrecht Germany). An overnight broth culture of bacteria was made in 5.0 ml of TSB, pelleted and resuspended and the bacterial density was then adjusted to ∼0.3 (OD_600_). In a separate tube, 5.0 ml of RPMI 1640+10% FBS (without antibiotics) was aliquoted, and 150 μl of the diluted bacteria (OD_600_ = 0.3) was added to prepare the working bacterial culture. The cell culture media was replaced with RPMI 1640+10% FBS media with and without bacterial suspensions and cells incubated for 2 hours at 37°C, 5% CO_2_ incubator. The wells were again washed and then incubated with 400 μl of 0.025% Triton X- 100 by pipetting, transferred into corresponding Eppendorf tubes and mixed by vortexing for 30 sec. The recovered bacteria were serially diluted in sterile PBS (up to 10^-4^), and 20 μl was spotted in TSA for CFU estimation. The plates were dried and incubated at 37°C along with previous plates containing serially diluted working bacterial culture spots. The next day, the colonies in each plate were counted. The relative adhesion was calculated using the following formulae:

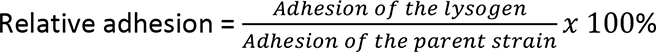

### 2.8 Proteomics of the secretome

The proteomics of the secretome was analyzed using a data-independent acquisition mass spectrometry (DIA-MS) using Orbitrap Fusion Lumos Tribrid Mass Spectrometer (Thermo Fisher Scientific, USA) with Dionex UltiMate™ 3000 UHPLC system (Thermo Fisher Scientific, USA) at The Flinders Omics Facility, Flinders University according to established protocol <u>(Supplementary</u> <u>text ST2)</u>. Briefly, a few colonies from overnight cultured NA plates were dissolved in sterile saline to obtain 1.0 MFU standard. One hundred microliters of the solution were added to 15.0 ml of TSB in a 50.0 ml Falcon tube and incubated at 37°C in a shaking incubator (180 rpm, 45° angle). After ∼7 hours, the tube was briefly vortexed and centrifuged at 4000 x g for 10 min. The secretome was sterilized by passing it through a 0.2 μm syringe filter (25 mm, Acordisc^®^, Pall International, Fribourg, Switzerland) and concentrated using a Pierce™ Protein Concentrator PES (3K MWCO, #Cat: 88525, Thermo Scientific, USA) by centrifuging the sample down to approximately 2-3 ml. The protein concentration was determined using NanoOrange™ Protein Quantitation Kit (Invitrogen, USA. #Cat: N6666) as per the manufacturer’s instruction. The proteins were then processed at the Flinders Omics Facility, Flinders University. The complete set-up and protocols are elaborated in <u>Supplementary text ST2.</u>

The DIA spectra were then processed and quantified using Spectronaut v15 (Biognosis AG, Schlieren, Switzerland) with default settings. A *S. aureus* proteome database created from all genes identified in SA333 was used as a reference. Gene annotations were assigned based on SA333 strain using Bakta v1.6.1 (Schwengers et al. 2021b). Differential protein expression analysis was performed in R v4.2.0 (https://www.R-project.org/) using the DEP package v1.20.0 to calculate differentially expressed proteins (DEP) (Zhang et al. 2018). The threshold for identifying differentially expressed proteins was set at a false discovery rate (FDR) of less than 0.05.

### 2.9 Bioinformatics and statistical analysis

Pairwise alignment and multiple-sequence alignment between the genomes were performed using MAFFT, and the phylogenetic tree was inferred using PhyML tree in Geneious Prime 2022.2.2 (http://www.geneious.com). The average nucleotide identity (ANI) of assembled genomes was calculated using a FastANI web tool available at https://proksee.ca (Jain et al. 2018). Statistical analysis was performed using Prism v9.4 (GraphPad Software, USA), and graphs were made in R v4.2.0 (https://www.R-project.org/). The difference in means of biofilm and metabolic activity was tested using the student t-test. P<0.05 was considered statistically significant unless stated otherwise.

### 2.10 Data availability

The genomic data (reads and assemblies) can be found in the Sequence Read Archive (SRA) under project number PRJNA914892, specifically with sample numbers SAMN32360844 (SA222) and SAMN32360890 (SA333).

## 3. RESULTS

### 3.1 Genomic features of SA222, SA333, lysogens and the prophages

Sequencing and genomic analysis revealed that there were minimal genetic variations between SA222 and SA333, which were isolated from the same CRS patient 567 days apart. The strains had the same 32.8% GC content, clonal complex (CC22) and sequence type (ST22) (**Figure 1A**; **Table 1**). The average nucleotide identity (ANI) between SA222 and SA333 was 99.99%, and the average aligned length between the two genomes was 2,390,601 bp (**Figure 1B**). The alignment also identified an additional φSa3int-group prophage (hereafter φSa3int prophage) in SA333. Similarly, the ANI between SA222 (the recipient strain) and lysogenized SA222 which had been genetically modified by lysogenization with φSa3int prophage (SA-L1, SA-L2) was ∼100%, with an average aligned length of 2,746,692 bp and the ANI between donor SA333 and lysogens (SA-L1, SA-L2) was 99.99%, with an average aligned length of 2,385,728 bp (**Figure 1B**). The key difference between the clinical isolates was that while SA222 harbored one intact prophage (φSa2int, 52,500 bp), SA333 had two intact prophages (φSa2int; 50,792 bp and φSa3int; 43,795 bp) (**Table 1**). Upon gene annotation, we could see that although the φSa2int prophage in SA333 lost some DNA compared to SA222, SA333 had gained two transposases (length = 1236 bp and 768 bp) **(Figure S1-A-D**). The identified φSa2int prophage in SA222 and SA333 was most closely related to Staphylococcus phage phi2958PVL (NC_011344, length = 47,342 bp), while the additional prophage in SA333 was most closely related to Staphylococcus phage IME1361_01 (NC_048657, length = 43,516 bp) (**Table 1**). There was only 60,421 bp of non-identical nucleotide bases between SA222 and SA333, including the 43,795 bp φSa3int prophage (**Figure 1D**), confirming that SA222 and SA333 were the same strain, but had gained the flexible prophage (φSa3int) at some point in time. Phylogenetic analysis and non-identical nucleotide differences between clinical isolates and the lysogens implied, as expected, that the laboratory-generated double lysogens were closer to SA333 than SA222 (**Figure 1C, 1D**). Surprisingly, the non-identical nucleotide bases between the recipient SA222 and SA-L1/L2 were 47,747 bp, which was 3,952 bp larger than the exact prophage identified suggesting some auxiliary cargo DNA and rearrangements in the lysogens during prophage integration (**Figure 1D**).

**Figure 1.**
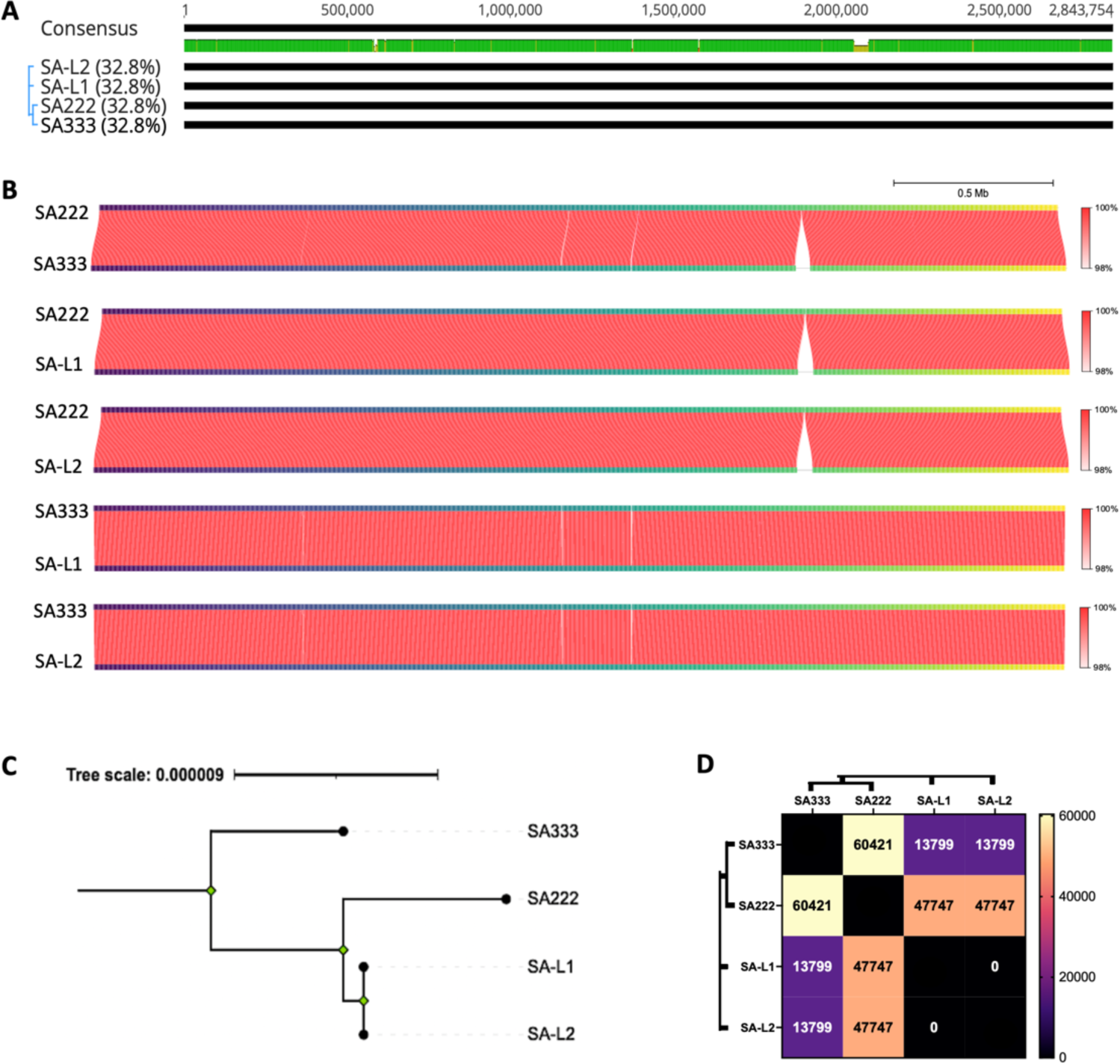
Multiple sequence alignments of clinical isolates and the lysogens. **(A)** Multiple sequence alignments of SA222, SA333 and laboratory generated double lysogens (two isolates; SA-L1 and SA-L2) show almost identical genome maps except for the prophage insert around 2.1 Mbp. **(B)** The average nucleotide identity (ANI) score between the clinical isolates and the lysogens further confirms that the prophage inserts on SA333 are the only significant genome change that has happened over time. The red colour gradient indicates the ANI percentage. **(C)** A rooted phylogenetic analysis between clinical strains and laboratory generated double lysogens reveals that as expected the double lysogens are genetically closer to the recipient strain SA222. **(D)** Distance matrix indicating the number of bases which are not identical.

**Table 1.**
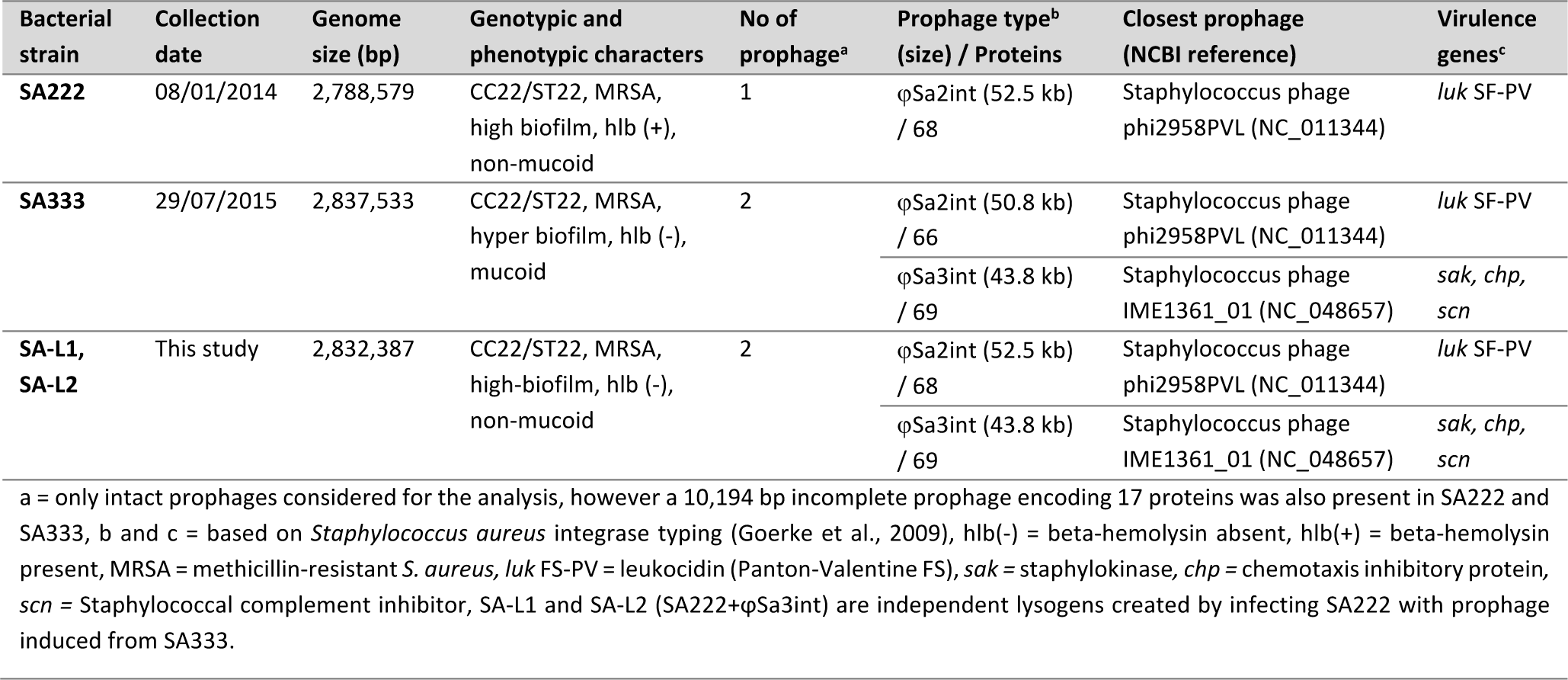
Details of the *Staphylococcus aureus* clinical isolates used in this study

| Bacterial strain | Collection date | Genome size (bp) | Genotypic and phenotypic characters | No of prophage<sup>a</sup> | Prophage type<sup>b</sup> (size) / Proteins | Closest prophage (NCBI reference) | Virulence genes<sup>c</sup> |
| --- | --- | --- | --- | --- | --- | --- | --- |
| SA222 | 08/01/2014 | 2,788,579 | CC22/ST22, MRSA, high biofilm, hlb (+), non-mucoid | 1 | φSa2int (52.5 kb) / 68 | Staphylococcus phage phi2958PVL (NC_011344) | luk SF-PV |
| SA333 | 29/07/2015 | 2,837,533 | CC22/ST22, MRSA, hyper biofilm, hlb (-), mucoid | 2 | φSa2int (50.8 kb) / 66<br>φSa3int (43.8 kb) / 69 | Staphylococcus phage phi2958PVL (NC_011344)<br>Staphylococcus phage IME1361_01 (NC_048657) | luk SF-PV<br>sak, chp, scn |
| SA-L1, SA-L2 | This study | 2,832,387 | CC22/ST22, MRSA, high-biofilm, hlb (-), non-mucoid | 2 | φSa2int (52.5 kb) / 68<br>φSa3int (43.8 kb) / 69 | Staphylococcus phage phi2958PVL (NC_011344)<br>Staphylococcus phage IME1361_01 (NC_048657) | luk SF-PV<br>sak, chp, scn |

a = only intact prophages considered for the analysis, however a 10,194 bp incomplete prophage encoding 17 proteins was also present in SA222 and SA333, b and c = based on *Staphylococcus aureus* integrase typing (Goerke et al., 2009), hlb(-) = beta-hemolysin absent, hlb(+) = beta-hemolysin present, MRSA = methicillin-resistant *S. aureus*, luk FS-PV = leukocidin (Panton-Valentine FS), sak = staphylokinase, chp = chemotaxis inhibitory protein, scn = Staphylococcal complement inhibitor, SA-L1 and SA-L2 (SA222+φSa3int) are independent lysogens created by infecting SA222 with prophage induced from SA333.

### 3.1 Prophages induced from clinical isolates and the lysogen displayed a multiple host-range

The prophage genome annotation revealed that φSa2int prophage mostly encoded phage-related genes and hypothetical genes (**Figure 2A**). Similarly, the φSa3int prophage encoded a complete set of IEC genes (*sak, chp, scn*) and phage-related genes (**Figure 2B**). Neither prophage encoded any antibiotic-resistance genes (ARGs) within their genomes. However, both prophages had a *clpP* gene encoding Clp protease involved in lysogenic-lytic switching (Thabet, Penadés and Haag 2022), indicating that both prophages should be capable of productive prophage induction (**Figure 2A-B**).

**Figure 2.**
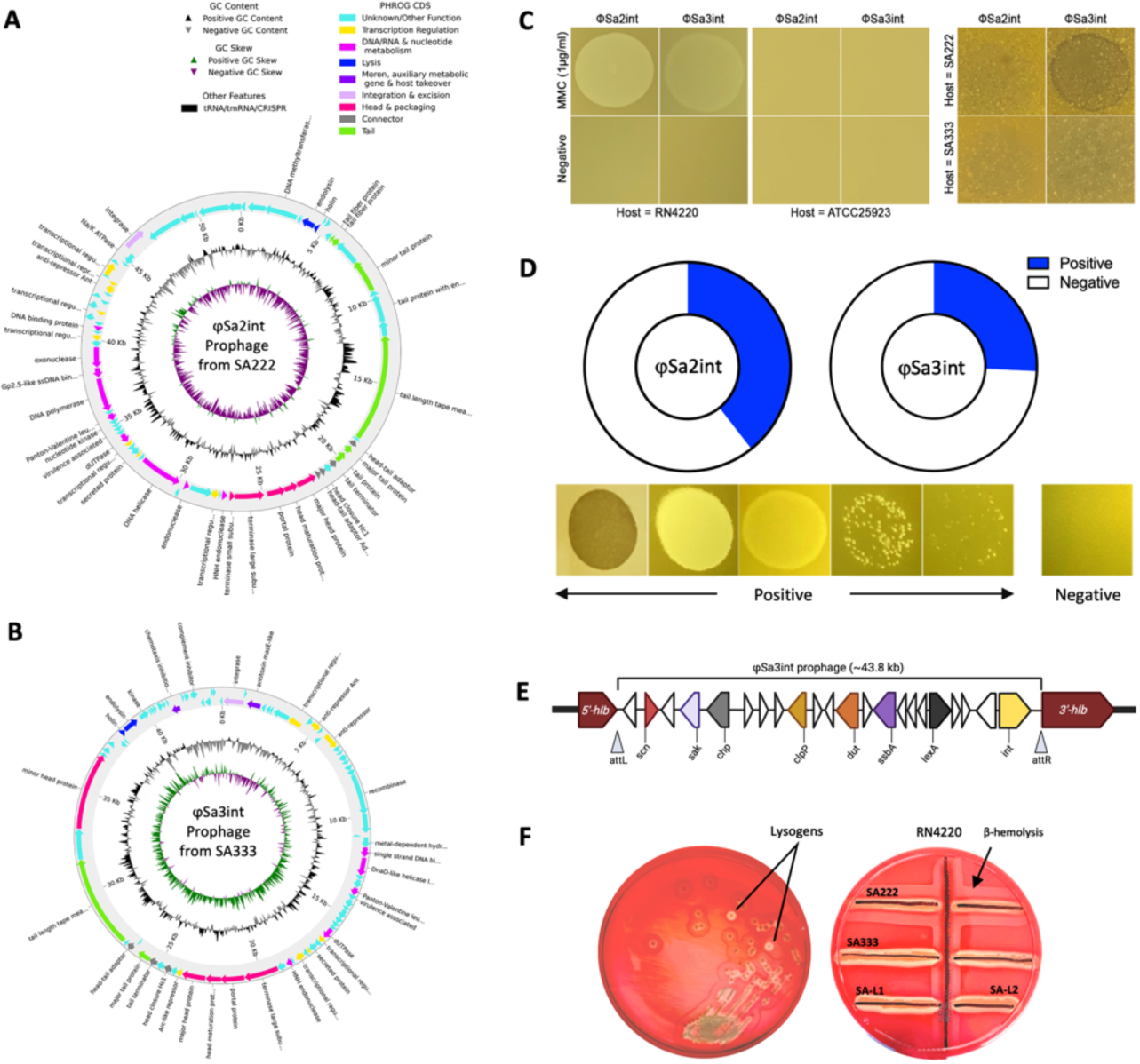
Annotation of identified prophages in SA222 and SA333 and their inducibility. **(A)** The φSa2int prophage is present in both clinical strains SA222 and SA333. The 52.5 kb prophage region primarily consists of hypothetical genes of bacterial as well as phage origin whose functions are as yet unknown. **(B)** The φSa3int prophage was present in clinical isolate SA333 in addition to φSa2int. The ∼43.8 kb prophage encoded major human immune evasion genes (*sak, scn, chp*). These genes are clustered together near the integrase (*xerC*). **(C)** Productive induction of prophage was observed from both clinical isolates (SA222 and SA333) using mitomycin (1.0 μg/ml). The induced phages showed clear lysis on indicator host RN4220 but did not show lysis on *S. aureus* ATCC25923. Further, the induced phage from SA333 (φSa3int) was able to lyse SA222, but the induced phage from SA222 (φSa2int) was unable to infect SA333 or SA222, possibly because of superinfection immunity. **(D)** The multiple host-range of induced phages from SA222 (φSa2int) and SA333 (φSa3int) indicated that the phages can infect and kill multiple clinical isolates. The φSa2int prophage from SA222 was able to infect almost 40% (26/66) clinical strains while φSa3int induced phage from SA333 could infect almost 26% (17/66) clinical isolates. The lower figure panels indicate the spots that were considered positive and negative. **(E)** Schematic chromosomal location of transduced φSa3int prophage and identified genes in SA-L1. The 43.8 kb insert was integrated within the *hlb* gene thereby truncating it. **(F)** Assessment of beta-hemolytic activity on sheep blood agar. The modified lysogens (SA-L1 and SA-L2) lost their β-hemolytic property.

Productive prophage induction was observed in both clinical strains, as lysis zones (plaques) were observed in the spot assay. A clear lysis zone was observed on indicator strain RN4220, indicating an intact lysogenic-to-lytic switch in the prophages, a productive induction, and a re-infectivity of φSa2int and φSa2int and/or φSa3int prophages induced from SA222 and SA333, respectively (**Figure 2C**). While induced prophage from SA333 (φSa2int and/or φSa3int) was also able to infect and lyse SA222 (lacks φSa3int prophage), induced prophage from SA222 (φSa2int) could not infect and lyse SA333 (which already has both φSa2int and φSa3int prophages), likely because of superinfection exclusion (**Figure 2C**). Similarly, induced prophage from lysogens (SA-L1 and SA-L2) could infect the parent strain SA222, indicating productive induction of transduced φSa3int prophage.

Further, released phages from both clinical isolates (SA222 and SA333) produced a partial or complete lysis spot on multiple clinical isolates using a spot assay, indicating a broad-host-range of the induced prophages and their ability to transduce the prophage-encoded virulence factors to multiple clinical isolates. Representative negative and positive lysis spots are shown in **(Figure D)**. Prophages induced from SA222 (φSa2int) could infect 39.4 % (26/66) of clinical strains, while productive prophages from SA333 (φSa2int and φSa3int) could infect 25.8 % (17/66) of clinical strains (**Figure 2D**). Phages released from both clinical isolates did not produce either a partial or a complete lysis spot on ATCC25923 (**Figure 2C**).

### 3.2 Integration of φSa3int prophage into *hlb* gene inhibited the production of β-hemolysin

Whole genome sequencing of the lysogens (SA-L1 and SA-L2) confirmed that a ∼43.8 kb φSa3int prophage DNA induced from SA333 was integrated into the SA222 chromosome within the *hlb* gene (start = 2,041,825 bp, end = 2,088,955 bp) **(Figure S1),** generating a double-lysogen with quadruple conversion (negatively converted *hlb* gene and incorporation of all three IEC genes: *sak, scn* and *chp*) (**Figure 2E**). The *hlb* gene was truncated near the 5’ end resulting in two incomplete genes; 201 bp (66 aa, molecular weight = 7321.09 amu) and 825 bp (274 aa, molecular weight = 31257.36 amu) **(Figure S2).** As such, the lysogens lacked β-hemolytic activity in sheep blood agar (**Figure 2F**). Genomic analysis also revealed that the φSa3int prophage in this study lacked *sea/sep* genes, which are commonly found prophage-encoded virulence factors in Sa3int-group prophages (**Figure 2B**).

### 3.3 Domestication of a φSa3int prophage had no impact on bacterial growth, biofilm biomass, metabolic activity and adhesion to primary human nasal epithelial cells

The domestication of the additional ∼43.8 kb ϕSa3int prophage DNA did not alter the growth kinetics of the lysogens compared to the recipient host SA222 (**Figure 3A**). Also, there was no significant change in biofilm biomass between the lysogens and SA222 (**Figure 3B**). The colony morphology of SA222 and SA-L1/SA-L2 on Congo red agar were similar. In contrast, the colony morphology of donor SA333 on Congo red media was wrinkled, suggesting strong mucoid production (**Figure 3C**). Further, the metabolic activity of the biofilm was similar between lysogens and recipient host SA222 (**Figure 3D**). However, the biofilm biomass and metabolic activity of SA333 (donor) were significantly higher than the laboratory-generated double lysogens, despite having a non-aligned nucleotide difference of 13,797 bp only (**Figure 1B, 1D**). Further, there was no significant difference in adhesion properties between the recipient SA222 and the double lysogens in primary HNECs. The relative adhesion of the lysogen to the HNEC compared to the recipient host SA222 was 94%, implying a slight reduction in the adhesion of *S. aureus* after infection with φSa3int prophage but did not reach statistical significance.

**Figure 3.**
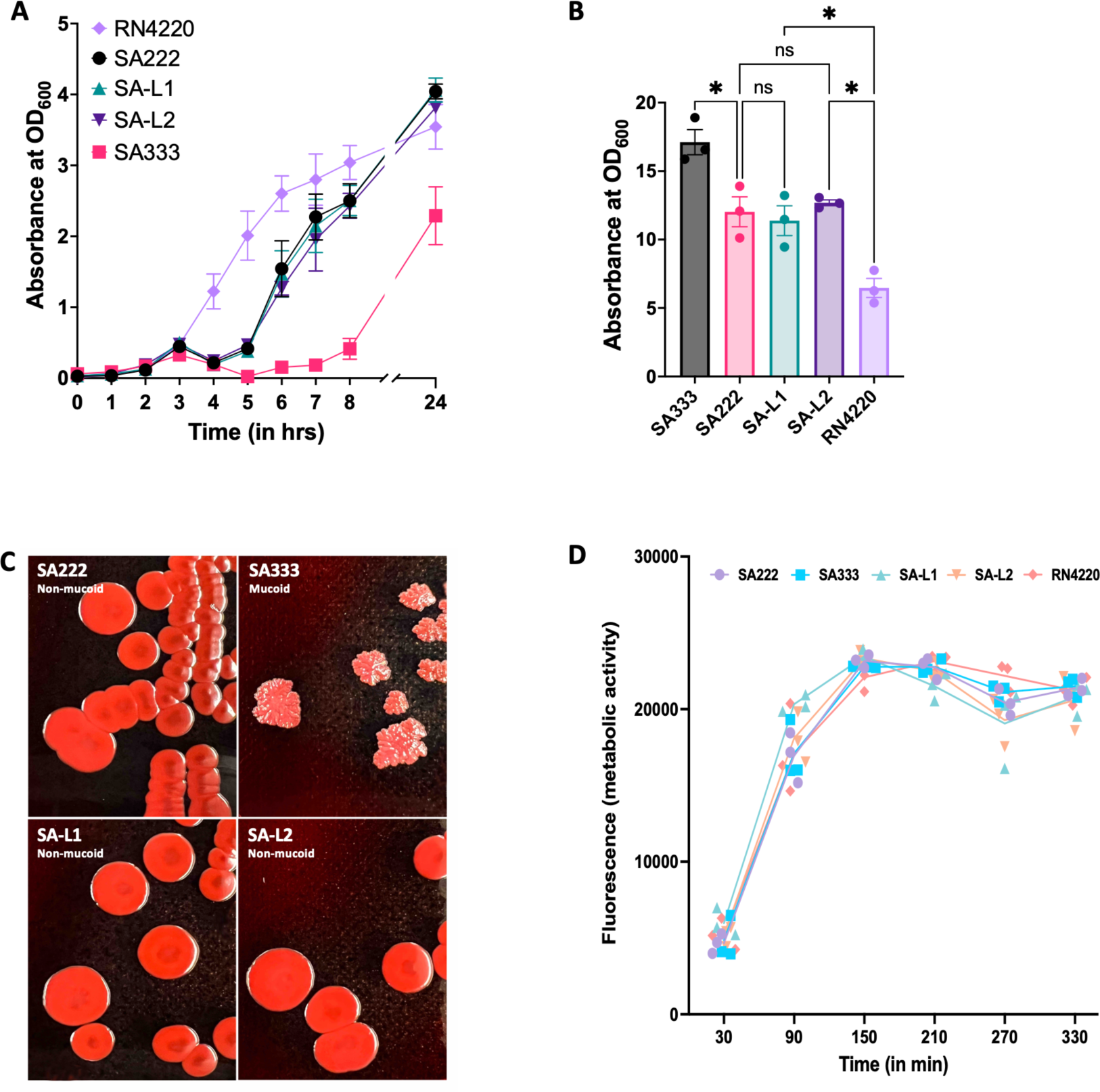
Comparisons of *in vitro* characteristics between SA222, SA333 and laboratory generated double lysogens (SA-L1 and SA-L2). **(A)** Bacterial growth curve in tryptic soy broth did not show any significant change in growth kinetics between recipient SA222 and double lysogens (SA-L1 and SA-L2), indicating domestication of an additional ∼47.1 kb ϕSa3int prophage did not impact the growth of the recipient strain SA222. **(B)** Biofilm estimation by crystal violet assay also indicated that the domestication of ∼47.1 kb ϕSa3int prophage did not impact biofilm formation. **(C)** Overnight culture of *S. aureus* strains in Congo red for qualitative estimation of mucoid phenotype further indicated that the ϕSa3int prophage and the genes it carried had no impact on phenotypic differentiation between mucoid and non-mucoid phenotype. **(D)** Study of biofilm metabolic activity by Alamar Blue assay indicated that there was no significant change in metabolic activity between recipient isolate (SA222) and double lysogens (SA-L1 and SA-L2).

### 3.4 Integration of φSa3int prophage equips *S. aureus* with human immune evasion factors

The proteomics of the secretome collected from SA222, SA333 and one of the laboratory-generated double lysogens (SA-L1) indicated that acquisition of the φSa3int prophage arms *S. aureus* bacteria with multiple prophage-associated human immune evasion factors. These include staphylokinase, SCIN, CHIPS and recombinase protein (recT) that is secreted as an exoprotein (**Figure 4A, cluster 1)**. Altogether, the lysogen significantly regulated thirty-eight exoproteins that were differentially expressed in the lysogen compared to its recipient strain SA222. Among them, twenty-one (55.3 %) were up-regulated including staphylokinase (sak), SCIN (scn), and intercellular adhesion protein B (icaB). In contrast, seventeen proteins (44.7 %) were down-regulated including β-hemolysin (hlb/sph) and outer membrane porin (phoE) (**Figure 4B-C**). Among these genes, 60.5% (23/38) were of unknown function. Similarly, despite having few genetic variations between lysogen and its donor host SA333 (**Figure 1B, 1D**), forty proteins were significantly differentially regulated (**Figure 4A, clusters 2 and 3)**. Among them, twenty-seven proteins (67.5 %), including enterotoxins SEO and SEG were up-regulated. In contrast, thirteen proteins (32.5 %), including elastin binding protein (ebp) were down-regulated (**Figure 4D-E**). As predicted, most of these proteins were of bacterial origin, and the hypothetical proteins, predominantly prophage encoded, constituted only 15.0% (6/40) of proteins because both strains (SA333 and SA-L1) carry identical mobile genetic elements (**Figure 1B**).

**Figure 4.**
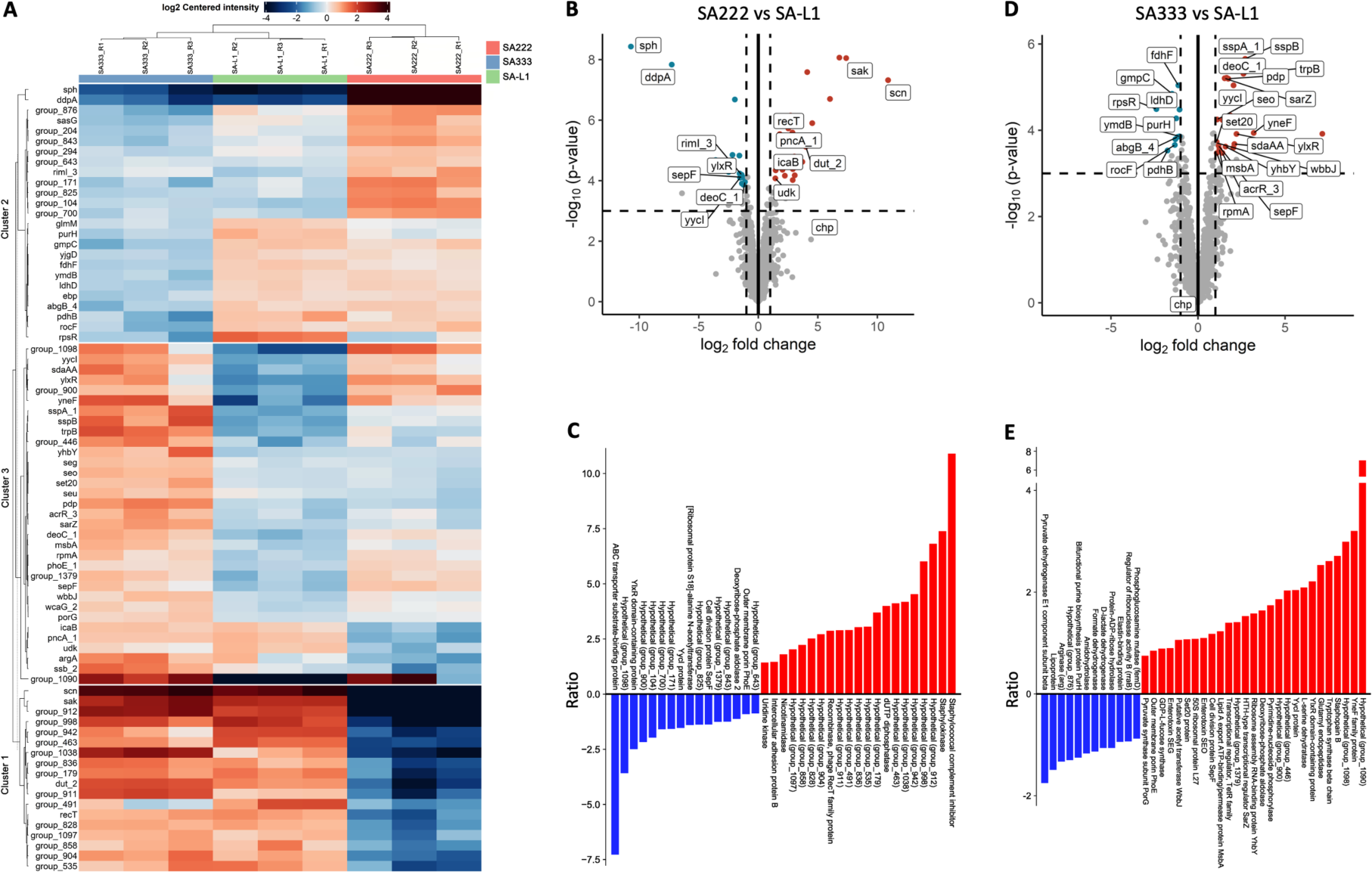
Proteomics of the secretome from SA222, SA333 and SA-L1. **(A)** The heatmap showing the differential expression between donor SA333, recipient SA222 and laboratory-generated double lysogen (SA-L1). The proteins in cluster 1 (including *sak, scn, recT, dut*) were significantly upregulated in SA-L1 compared to recipient isolate SA222. The expression of genes in this group was similar between the lysogen and the donor isolate SA333. **(B)** Volcano plot showing the differential expression of proteins between recipient host SA222 and lysogen (SA-L1). **(C)** A bar plot showing the relative ratio of significantly up-and down-regulated proteins between SA222 and SA-L1. The virulence factors responsible for human immune evasion (sak and SCIN) are significantly upregulated along with the intercellular adhesion protein B (icaB). **(D)** Volcano plot showing the differential expression of proteins between donor host SA333 and lysogen (SA-L1). **(E)** A bar plot showing the relative ratio of significantly up-and down-regulated proteins between SA333 and SA-L1. There was no significant change in virulence factors encoded by prophage φSa3int, indicating the difference in phenotypes is not associated with prophage-associated genes or functions. P < 0.05 was considered significant in all the above analyses.

## 4. DISCUSSION

The productive induction of prophages from a lysogen leads to the release of phage particles that can either kill the competing strains or lysogenize the susceptible strains (Matos et al. 2013). It is now established that various drugs, antibiotics and even dietary compounds enhance prophage induction and promote the lysogenic conversion (Boling et al. 2020, Sutcliffe et al. 2021, Goerke et al. 2006). The prophages can also be induced spontaneously in response to nutrient availability and/or other environmental conditions, including niche variation. As over 90% of human clinical isolates carry Sa3int-group prophages, predominantly integrated into the *hlb* gene (van Wamel et al. 2006), we aimed to elucidate the contribution of φSa3int prophage domestication in *S. aureus,* particularly in the secretome of the bacteria. To achieve this, we introduced a φSa3int prophage (also called β-hemolysis-converting prophage) induced from a patient-derived double-lysogen (SA333), into a genetically similar single (φSa2int)-lysogen (SA222), isolated from the same patient almost 2 years earlier, to create a laboratory-generated double-lysogen. We then studied its phenotypic characteristics as well as proteomics. We chose these clinical isolates for our experiments because, our preliminary study suggested that although there were significant phenotypic variations between SA222 (high-biofilm, hlb(+)) and SA333 (hyper-biofilm, hlb(-)), the two CIs were genetically similar except for the gain of φSa3int prophage by SA333 and a few SNPs (data not shown) and also were collected from the same patient who had severe CRS.

Lysogenic conversion in *S. aureus* arms bacteria with many survival fitness traits like virulence factors, toxins, and biofilm upregulation and is common in clinical strains (Fernandez et al. 2018, Naorem et al. 2021, Bae et al. 2006). Prophage-mediated enhancement of biofilm has been observed in various bacteria like *Enterococcus faecalis* (Rossmann et al. 2015), *Streptococcus pneumoniae* (Carrolo et al. 2010), *Escherichia coli* (Li et al. 2022) and *S. aureus* (Fernandez et al. 2018). Further, multiple studies have described prophage-mediated phenotypic alteration, host adaptation and pathogenesis (Liu et al. 2020, Resch et al. 2013, Bobay, Rocha and Touchon 2013).

Our findings support the notion that lysogenic conversion in clinical isolates is common (Chaguza et al. 2022), as almost 40% of the CIs tested could be infected with a productive prophage derived from another clinical isolate, meaning they could gain prophage-associated virulence and pathogenicity via phage infection. Further, the variations in the strength of lysis spots in multiple host-range assays indicated that the induced temperate phages had varying degrees of susceptibility among CIs, suggesting variable lysogenic conversion of the host.

Several studies on *S. aureus* clinical isolates have suspected Hlb as an important non-pore-forming toxin promoting colonization and impacting ciliary clearance of bacteria in an animal model (Katayama et al. 2013, Seop Kim et al. 2000). A similar function of Hlb has thus been speculated in humans. However, most human-associated *S. aureus* lack β-hemolysin because of the integration of the Sa3int-group prophage in the *hlb* gene (van Wamel et al. 2006, Nepal et al. 2021). A similar truncation of the *hlb* gene in this study is seen after integrating φSa3int prophage DNA, thereby completely disrupting the production of the Hlb protein in the secretome. This seems counterintuitive as Hlb is a strong sphingomyelinase that promotes sphingomyelin degradation and stimulates biofilm formation in the presence of eDNA (Huseby et al. 2007, Huseby et al. 2010). Thus, we speculate that nasal colonizers might trade-off the Hlb function by gaining the immune evasion factors that are encoded by the φSa3int prophage and are known to protect bacteria from phagocytosis. However, re-expression of Hlb has been noted in *S. aureus* isolated from cystic fibrosis patients upon antibiotic (ciprofloxacin or trimethoprim) treatment and increased frequency of genomic alterations have been associated with prophage mobilization (Goerke et al. 2004). Other studies have shown the conditional excision of φSa3int prophage in a sub-set of the population in *in vivo* conditions and *S. aureus* thriving as a heterogeneous population that aggravates the infection (Guan et al. 2021). As the laboratory generated φSa3int prophage in our study could be re-induced, we support the notion that φSa3int prophage integration and excision are conditional and largely depend on external factors. However, more *in vivo* study is required to confirm the hypothesis and understand the conditional switching of *S. aureus* from Hlb positive to negative or vice versa.

*S. aureus* biofilm and its adhesion to various surfaces are important features that aid bacteria in colonizing and establishing themselves in various niches. Fernandez et al. suggested the role of *S. aureus* prophages in biofilm development and adhesion (Fernandez et al. 2018). Other researchers have suggested that the eDNA released during spontaneous prophage induction acts as a quorum sensor leading to an enhanced biofilm formation (Carrolo et al. 2010). However, we did not observe significant enhancement of biofilm production or adhesion to HNECs when comparing SA222 and SA222 following introduction of the φSa3int prophage, although biofilm-associated intercellular adhesion protein (icaB) was upregulated in the double lysogen. The contradictory findings may be due to the different prophage type used for lysogenization because Fernandez et al. used φ11 (Sa5int-group) and φ80 (Sa6int-group) to lysogenize the lab strain RN4220, while we used a Sa3int-group prophage to lysogenize another clinical isolate. Further, the possible explanation for non-significant biofilm change may be because the recipient host SA222 was already a high-biofilm forming isolate, and the contribution of integrated prophage was minimal. Although the result was unexpected, the observation ruled out a major contribution of φSa3int prophage including all the hypothetical genes it carries in biofilm development and adhesion of *S. aureus*. Since *S. aureus* SA333 and lysogens were genetically almost identical (99.51 % similar) except for a few SNPs and insertion sequences (data not shown) but had different biofilm, adhesion and mucoid phenotypes, the result has helped us narrow down on other possible bacterial genes responsible for hyper-biofilm characteristics. However, further experiments with knock-out mutants are required to confirm the role of these candidate genes in biofilm development.

Further, our results suggest that although integration of a Sa3int-group prophage seems expensive in terms of replication energy cost and also disrupts the β-hemolysin expression, the prophage equips the host bacteria with a multitude of accessory virulence factors like *sak, scn, chp, icaB*. The high incidence of human-specific anti-innate immunity factors in *S. aureus* isolated from humans is well known (Rohmer and Wolz 2021). It is also established that φSa3int prophages carry and disseminate IEC genes (*sak, chp, scn, sea/entA*), either partial or a complete set (Rohmer and Wolz 2021). *In vitro* secretome profiling of the SA222, SA333 and lysogen (SA-L1) revealed major changes in virulence, particularly human immune evasion modulation of *S. aureus* associated with the domestication of the prophages. Our results confirm and extend the existing knowledge that φSa3int-group prophages carry IEC genes in a cluster that are significantly upregulated and secreted as exoproteins. Staphylokinase protein (Sak) encoded by *sak* gene is a potent plasminogen activator that converts plasminogen into plasmin. Sak-mediated plasmin activity increases the local invasiveness of *S. aureus* leading to skin disruption and reduced clearance of bacteria by the host (Peetermans et al. 2014). Chemotaxis inhibitory protein (CHIPS) encoded by *chp* gene counters the first line of host defence, specifically inhibiting the response of human neutrophils and monocytes to complement anaphylatoxin C5a and formylated peptides. It directly binds to the C5a and formylated peptide receptors, preventing phagocytosis of the bacterium (Postma et al. 2004). Similarly, Staphylococcal complement inhibitor protein (SCIN) encoded by *scn* gene also counters the first line of host defence. It efficiently inhibits opsonization, phagocytosis and the killing of *S. aureus* by human neutrophils (de Jong et al. 2018). All of these genes were significantly upregulated in the lysogen secretome along with downregulation of hemolysin, indicating a quadruple conversion of SA222 by φSa3int prophage induced from SA333. These results also hint that the patient isolate SA222 gained a φSa3int prophage to establish itself in the nasal niche, as multiple CIs isolated after this time point from the same patient were also double-lysogens with IEC genes (data not shown).

All these observations together indicate that despite an increase in the genome size of the double-lysogen by almost 43.8 kb of Sa3int-group prophage DNA, there was no significant change in growth kinetics, biofilm and adhesion properties of *S. aureus*. However, the proteomics analysis of the secretome clearly indicated that lysogenization by Sa3int-group prophage DNA armed the bacterial host with additional flexible virulence features likely to help bacteria evade the human immune system by avoiding phagocytosis. Further, as induced phages could infect other clinical isolates but not their own parental host, we can assume that prophage domestication not only increases the virulence of the lysogenized host but also can landscape the microbiome, which may lead to bacterial dysbiosis and ultimately pathological conditions. Temperate phages (which are formed by productive prophage induction) have been observed in various environments like human gut, chronic wounds, and cystic fibrosis patients aggravating the disease. As recent advances in genomic sequencing have revealed that most *S. aureus* clinical isolates adapted to humans’ harbour prophages in their genome, and not all of the clinical isolates carry similar mobile genetic elements, verification of prophage-encoded toxigenic trait is of paramount importance in inflammatory diseases like CRS.

## 5. CONCLUSION

We conclude that lysogenic conversion of *S. aureus* by φSa3int prophage alters the bacterium’s virulence by upregulating the human immune evasion factors like sak, scn and chp and downregulating beta-hemolysin, a sphingomyelinase. Our research further confirmed that the growth, biofilm and adhesion of *S. aureus* are not associated with Sa3int-group prophage domestication. These findings demonstrate the need to consider mobile genetic elements like prophages while developing a treatment strategy in chronic diseases like CRS, as strains with/or without prophages have different virulence properties and risks of chronic colonization, despite having almost identical core genomes.

### Limitations and future direction

Although we could affirm the origin of human immune evasion cluster genes in *S. aureus* and predict the location of φSa3int-group prophage insertion with accuracy, we could not identify the genes responsible for increased biofilm and adhesion in SA333. However, the successful introduction of φSa3int prophage into SA222 ruled out the role of prophage and prophage-associated genes in the high biofilm and mucoid phenotype of *S. aureus* SA333, redirecting future research to the limited number of SNPs and insertion sequences that are present on SA333 but not on the lysogens.

### Funding information

RN was supported by THRF-BHI Postgraduate Research Scholarship and The University of Adelaide Scholarship. GH was supported by The University of Adelaide International Scholarships and a THRF Postgraduate Top-up Scholarship. SV was supported by a senior fellowship from the Passe and Williams Foundation.

### CRediT author statement

Conceptualization: SV, RN; Methodology, formal analysis and investigation: RN, GH, GB, MR, GS, SF; Data curation: GH, GB; Writing – original draft: RN, GH, GB; Writing – review & editing: GH, GB, SF, MR, GS, KS, AJP, PJW, SV; Visualization: RN, GH, GB; Supervision: SV, PJW, AJP; Funding acquisition: SV, PJW.

### Conflicts of interest

The authors declare that the research was conducted in the absence of any commercial and/or financial relationships that could be construed as a potential conflict of interest.

## Acknowledgements

We’d like to extend our sincere thanks to all the members of ENT Surgery Research Group, Basil Hetzel Institute (BHI) for their constructive suggestions during the research. We’d also like to extend our sincere thanks to past/present members of the group who were directly or indirectly involved in the collection of the clinical data.

## Supplementary data

All the supplementary data, protocol and materials pertaining to this research is available in a public database under the following doi address: http://10.6084/m9.figshare.22696627

### Glossary

Active lysogen: bacterial strain harboring identifiable prophage sequence and releasing reinfecting phage particles.
Double lysogen: bacterial cell harbouring two complete/intact prophages
Lysogen: bacterial cell containing one or more prophages within its genome
Lysogenic conversion: phenotypic change in a host bacterium caused by the integration of a prophage.
Lysogeny: state of phage integration into the bacterial genome.
Non-lysogen: bacterial strains lacking any identifiable prophage sequence in their genome
Passive lysogen: bacterial strains harboring identifiable prophage sequence but not releasing actively reinfecting phage particles
Productive induction: excision of prophage from the bacterial chromosome followed by release of phage particles either spontaneously or under external stress
Prophage: temperate phage DNA integrated in the bacterial genome.
Single lysogen: bacterial cell harbouring only one complete/intact prophage
Temperate phage: bacterial virus that can integrate into a bacterial genome (or be maintained extra-chromosomally), become stabilized in this way, and, upon receiving a cue, can excise and propagate.
Transduction: process of horizontal gene transfer, wherein a region of a bacterial genome is packaged into phage particles that, upon release and entrance into a new host, is inserted into the genome of the latter.

## Supplementary figures and figure legends

**Figure S1.**
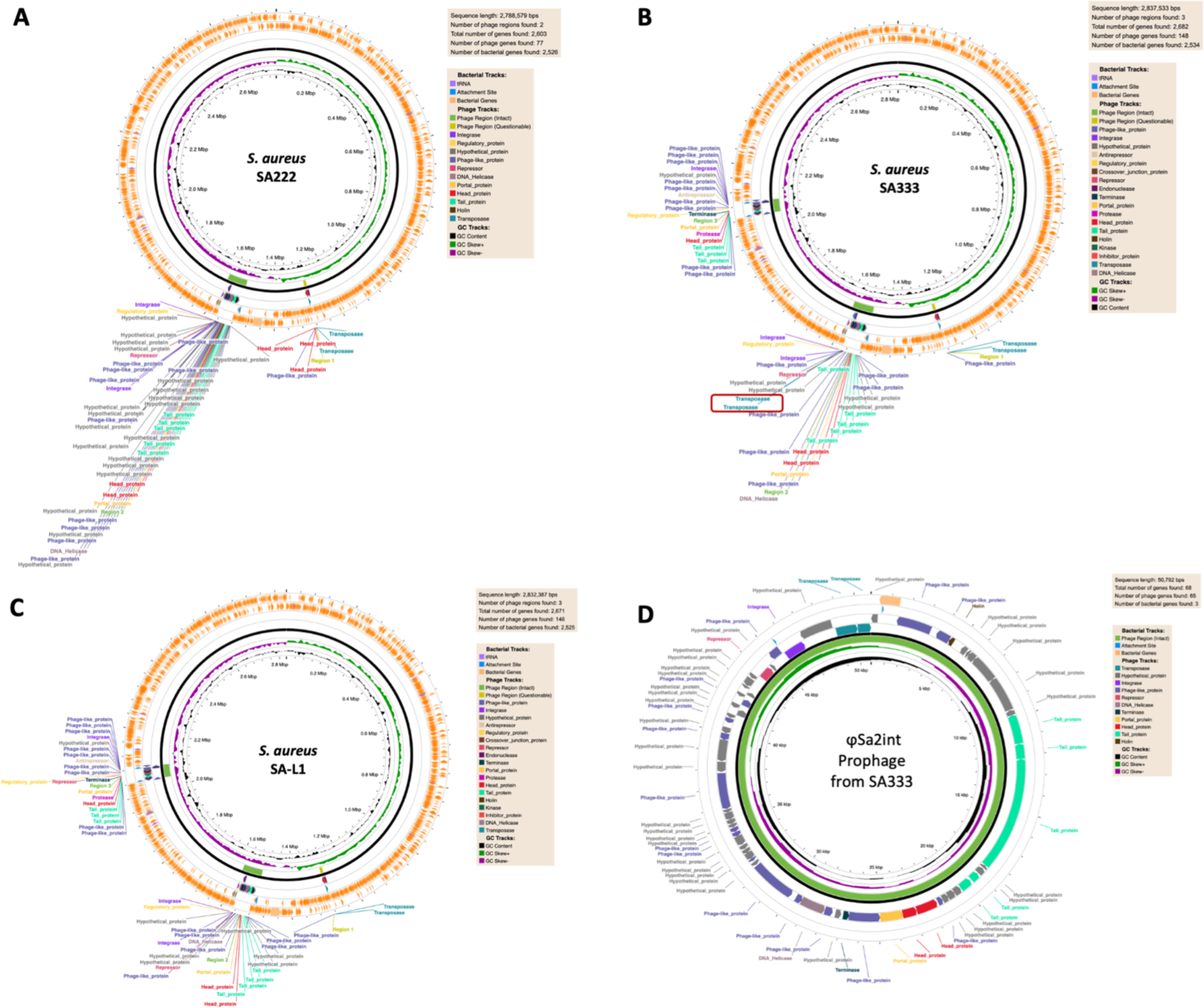
Identification of prophage region of SA222, SA333 and SA-L1 by PHASTEST. **(A)** *S. aureus* SA222 only harbours one intact prophage region (52.5 kb). **(B)** *S. aureus* SA333 harbours two intact prophage regions (50.8 kb and 43.8 kb). It is noted that the first prophage in *S. aureus* SA333 is almost similar to the one from *S. aureus* SA222 but has two transposases integrated into the prophage region (red box). **(C)** Laboratory-generated *S. aureus* SA-L1 harbours two intact prophages, one from *S. aureus* SA222 and one from *S. aureus* SA333. The second prophage was induced from *S. aureus* SA333 and inserted into *S. aureus* SA222. **(D)** The genetic mapping of φSa2int prophage from *S. aureus* SA333. Note the gain of two transposase enzymes compared to the same prophage present in *S. aureus* SA222.

**Figure S2.**
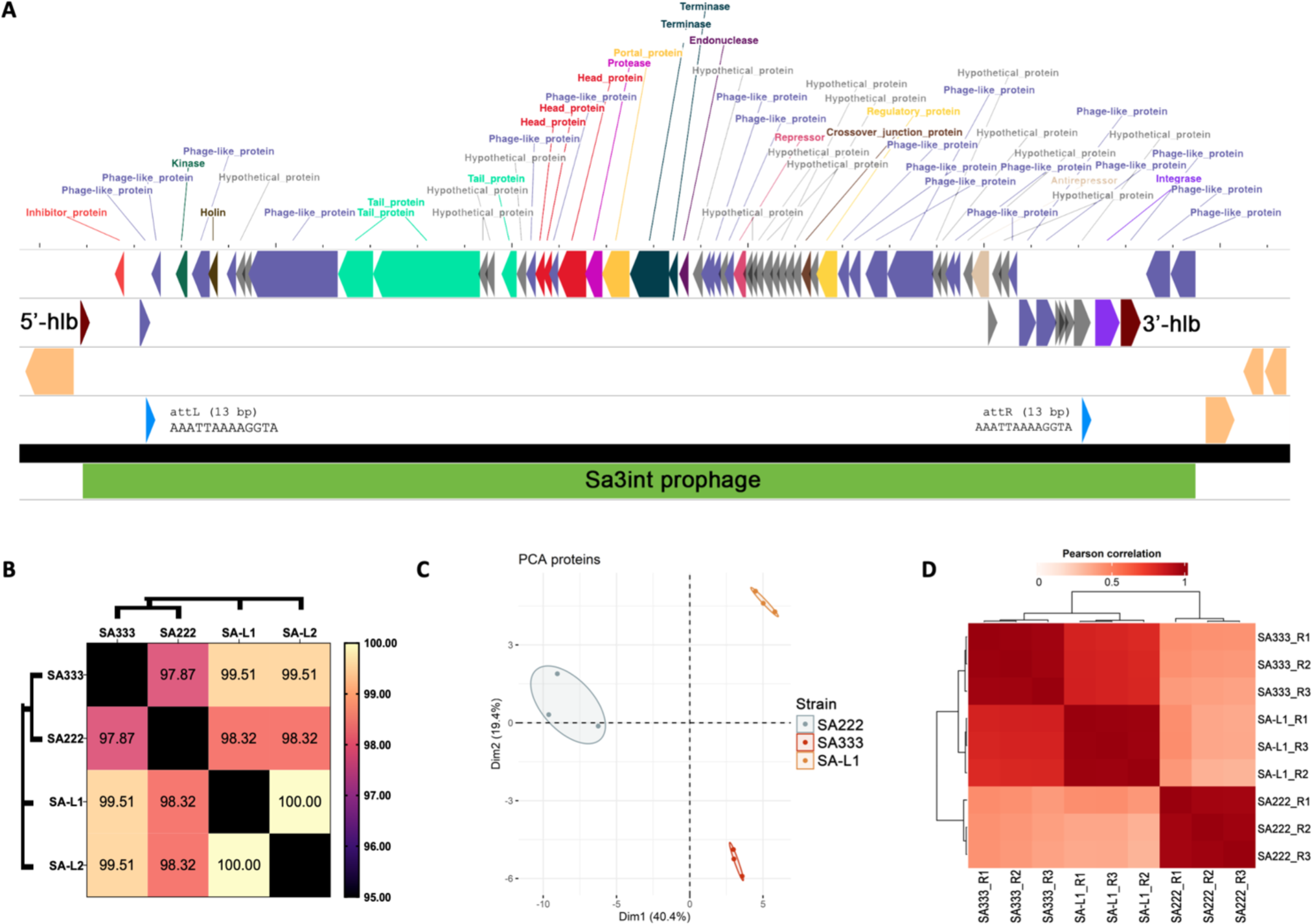
Chromosomal location φSa3int prophage and correlations between SA222, SA333 and SA-L1. **(A)** Chromosomal location of the φSa3int prophage insert in SA-L1. **(B)** A heatmap representing the genomic similarities between donor (SA333), recipient (SA222) and laboratory-generated double lysogen (SA-L1 and SA-L2). The numbers inside the square represent aligned identical bases/nucleotides between the strains in percentage. **(C)** Principal component analysis (PCA) of proteomics (triplicates) between SA222, SA333 and SA-L1. The analysis shows that the proteomics of triplicates clustered together, indicating consistency of the secretome. **(D)** Pearson’s correlation of proteomics (triplicates) between SA222, SA333 and SA-L1 also represents consistency in release factors in the secretome.

## Notes

http://10.6084/m9.figshare.22696627

### Competing Interest Statement

The authors have declared no competing interest.

